# inquiSTR: a toolkit for accurate and efficient population-scale tandem repeat genotyping and analysis

**DOI:** 10.64898/2026.06.09.731080

**Authors:** Wouter De Coster, Fahri Küçükali, Kristel Sleegers, Rosa Rademakers

**Author notes:** Corresponding Author Wouter De Coster, VIB Center for Molecular Neurology, Universiteitsplein 1, 2610 Wilrijk, Belgium.

## Abstract

Tandem repeats are highly mutable genomic elements linked to human traits and diseases. Profiling large catalogs of tandem repeats from population-scale long-read sequencing data requires accurate and efficient tools. We introduce inquiSTR, a command-line toolkit for fast genome-wide tandem repeat length genotyping. inquiSTR, with efficient parallel processing and low-memory streaming algorithms, genotypes a genome-wide repeat catalog of 1.78 million loci in less than two minutes. Benchmarking shows high accuracy and significantly faster performance compared to existing tools and truth sets. inquiSTR also provides methods for downstream analyses such as population structure inference, association testing, and outlier detection.

## Background

Tandem repeats (TRs) are direct head-to-tail repetitions of a DNA motif, and represent some of the most mutable elements of the human genome (Ibañez et al. 2024). TRs show length variation across individuals, populations, and somatically within an individual, and contribute to genome instability, gene regulation, traits, and disease susceptibility (Depienne and Mandel 2021). Genome-wide repeat catalogs, such as the adotto and TRexplorer catalogs, comprise approximately 1.8-5 million loci (Weisburd et al. 2025; English et al. 2024). Repeat expansions underlie numerous human disorders, such as Huntington’s disease and frontotemporal dementia/ALS, making the detection of expansions critical for both basic research and clinical applications (Renton et al. 2011; DeJesus-Hernandez et al. 2011; Gusella et al. 1983). Their repetitive nature and length heterogeneity typically require long-read sequencing for accurate resolution, and several tools have already been developed for TR calling on long-read sequencing data (Ziaei Jam et al. 2024; Dolzhenko et al. 2024; Lougheed et al. 2026; Aliyev et al. 2026; Chiu et al. 2021). However, faster yet accurate repeat-genotyping software is required to accommodate the increasingly large-scale long-read population sequencing initiatives. Furthermore, once population-scale TR genotypes are obtained, a range of downstream analyses remains, including visualization, association testing, and outlier analysis.

To address these challenges, we present inquiSTR (/In’kwizitər/, pronounced like “inquisitor”), a toolkit for rapid, genome-wide TR length genotyping from long-read sequencing data, with an architecture designed to scale to hundreds of samples and millions of TR loci, leveraging parallelization and streaming to improve computational efficiency. Through benchmarking, we demonstrate that inquiSTR substantially improves genotyping speed without sacrificing accuracy relative to existing long-read TR genotypers and works well in tandem with sequence-based TR genotypers. We also show that inquiSTR can genotype pathogenic expanded alleles and can be used to statistically identify disease- or trait-relevant expansions via association or outlier analysis. By facilitating large-scale, cost-effective TR profiling, inquiSTR enables systematic investigation of TR variation and its contribution to human traits and disease in population genomics.

## Results

inquiSTR demonstrated excellent performance and scalability, genotyping the Adotto catalog (used for the GIAB truth genotypes; 1.78M loci) in 103 seconds for PacBio data (2 threads) and 232 seconds for ONT data (6 threads). We observed a substantial speedup for ONT data when increasing parallelization from 1 to 4 threads, while running inquiSTR on PacBio data did not benefit from more than 2 threads (**Figure 1A**). Increasing the thread count further did not improve efficiency due to IO limitations, but these findings are likely compute architecture-dependent. The memory-use profile was flat up to 5 threads, after which it increased linearly (**Supplementary Figure 1**). When comparing inquiSTR to recently developed and well-performing TR genotypers, TRGT (Dolzhenko et al. 2024) on PacBio data and LongTR (Ziaei Jam et al. 2024) on ONT data, we demonstrate that inquiSTR genotyping is faster (respectively 8.66x and 76.6x) and more memory-efficient, and an approach with inquiSTR genotyping and filtering of TR loci before full sequence genotyping with TRGT or LongTR is beneficial for both time and memory use (**Figure 1B+C**; **Supplementary Figure 3**). When comparing inquiSTR to Straglr, another length-based genotyper, we observe that inquiSTR is 233x faster on ONT data and 488x faster on PacBio data (**Supplementary Figure 2**).

**Figure 1:**
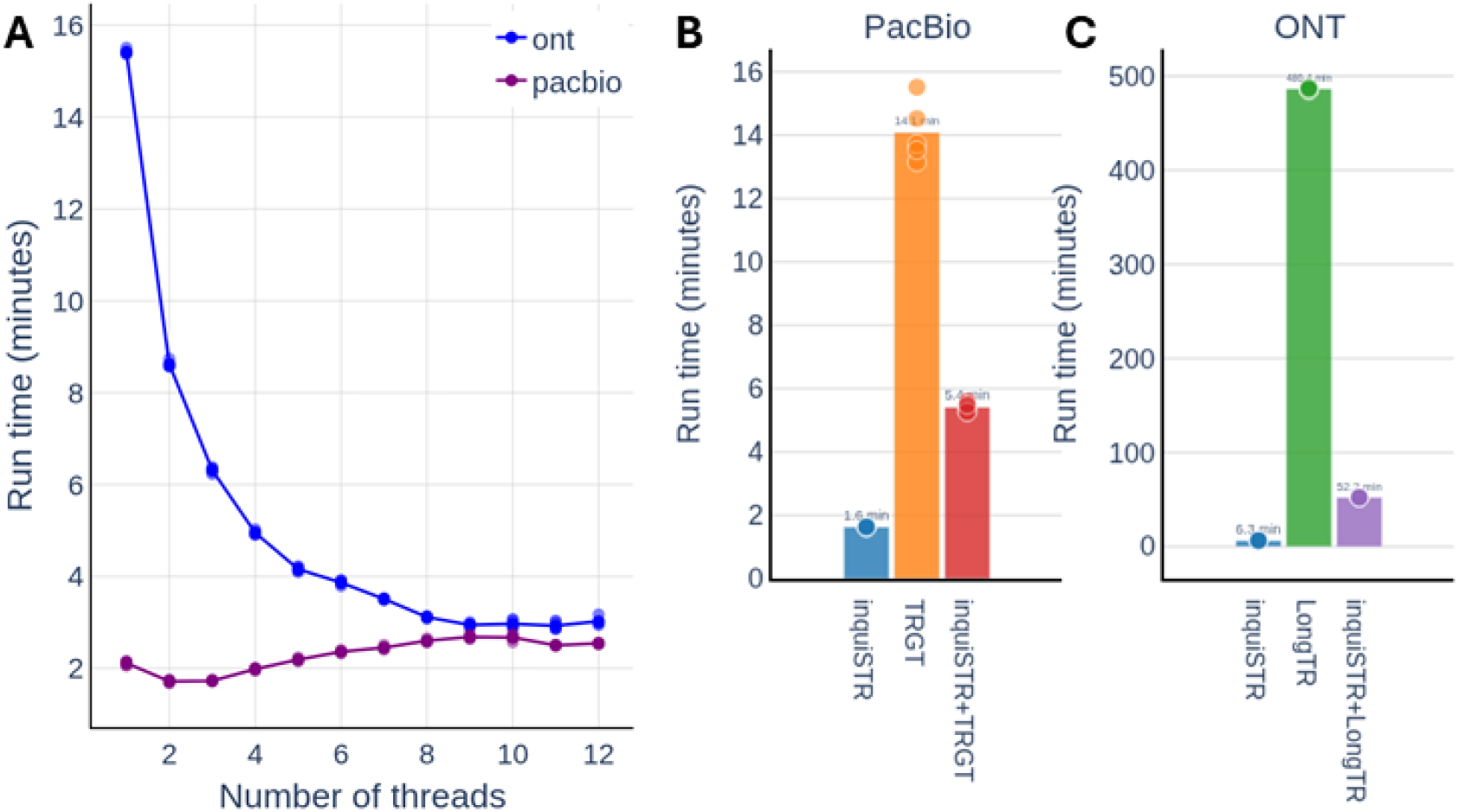
**A** performance of inquiSTR for genotyping the Adotto catalog (1.78M loci) with an increasing number of threads. **B+C**: comparison of run time for the PacBio (**B**) and ONT (**C**) dataset, showing inquiSTR call genotyping compared to TRGT and LongTR. Running inquiSTR in tandem with TRGT or LongTR and subsetting the loci of interest to those that show length variation before full ‘sequence-level’ genotyping is beneficial.

We next compared TR genotypers based on TR length accuracy (**Table 1**). When comparing inquiSTR genotypes from the PacBio dataset to the GIAB truth genotypes, we observe 99.83% of genotypes are within 3 bp of an exact match, with 94.54% being exact matches. For the ONT dataset, 99.65% of genotypes are within 3 bp of an exact match, with 92.13% being exact matches. When comparing the TR lengths from TRGT and LongTR against the GIAB truth genotypes, we observe that inquiSTR TR lengths match the truth at least as well as either tool. We also demonstrated high concordance between the inquiSTR genotypes and those obtained from TRGT and LongTR within a 3 bp tolerance distance, although lower percentages are observed for exact matches with TRGT and LongTR, suggesting systematic small and off-by-one differences without a considerable effect on the actual genotype.

**Table 1:** Accuracy of HG002 TR lengths from inquiSTR, TRGT, and LongTR.

| Technology | Test set | Truth set | Within 3 bp | Within 1 bp | Exact matches |
| --- | --- | --- | --- | --- | --- |
| PacBio | inquiSTR | GIAB truth | 99.83% | 99.16% | 94.54% |
| ONT | inquiSTR | GIAB truth | 99.65% | 97.28% | 92.13% |
| PacBio | TRGT | GIAB truth | 99.38% | 98.36% | 94.45% |
| PacBio | LongTR | GIAB truth | 99.41% | 98.25% | 90.53% |
| ONT | LongTR | GIAB truth | 97.71% | 89.51% | 72.50% |
| PacBio | inquiSTR | TRGT calls | 99.57% | 98.07% | 79.93% |
| PacBio | inquiSTR | LongTR calls | 99.63% | 97.29% | 76.26% |
| ONT | inquiSTR | LongTR | 97.10% | 84.93% | 60.60% |

Discordant genotypes between TRGT and inquiSTR are dominated by loci with large softclips. Some are bona fide repeat expansions, while others are other structural variants in repetitive regions, such as (pericentromeric) rearrangements. Using softclips increases sensitivity for long repeat alleles but may result in false-positive expansions when a rearrangement overlaps a TR locus. The --require-spanning flag of `the inquiSTR call` can be used to set TR alleles supported only by soft-clipped reads to missing (NaN), but this affected only 0.04% of the loci in the PacBio dataset and 0.01% in the ONT dataset, without changing the overall accuracy metrics.

We also demonstrated that inquiSTR can identify pathogenic expansions (**Figure 2A**), and that genotyping settings for greater sensitivity (--imbalance 0.1) do not yield false-positive long-expanded alleles. Evaluation of population structure using PCA was performed with a selection of polymorphic repeats, demonstrating separation of superpopulations (**Supplementary Figure 4**). In an in-house nanopore genome-sequencing cohort of patients with frontotemporal dementia, after running `inquiSTR call` with the Adotto catalog, the `inquiSTR outlier` command correctly prioritized the C9orf72 repeat, a known cause for frontotemporal dementia, in two known carriers. We also used the `inquiSTR association` subcommand to test the association for a recently identified repeat expansion in frontotemporal dementia (De Coster et al. 2026) (**Figure 2B**), for which we visualized the repeat lengths in patients and controls using `inquiSTR plot` (**Figure 2C**).

**Figure 2:**
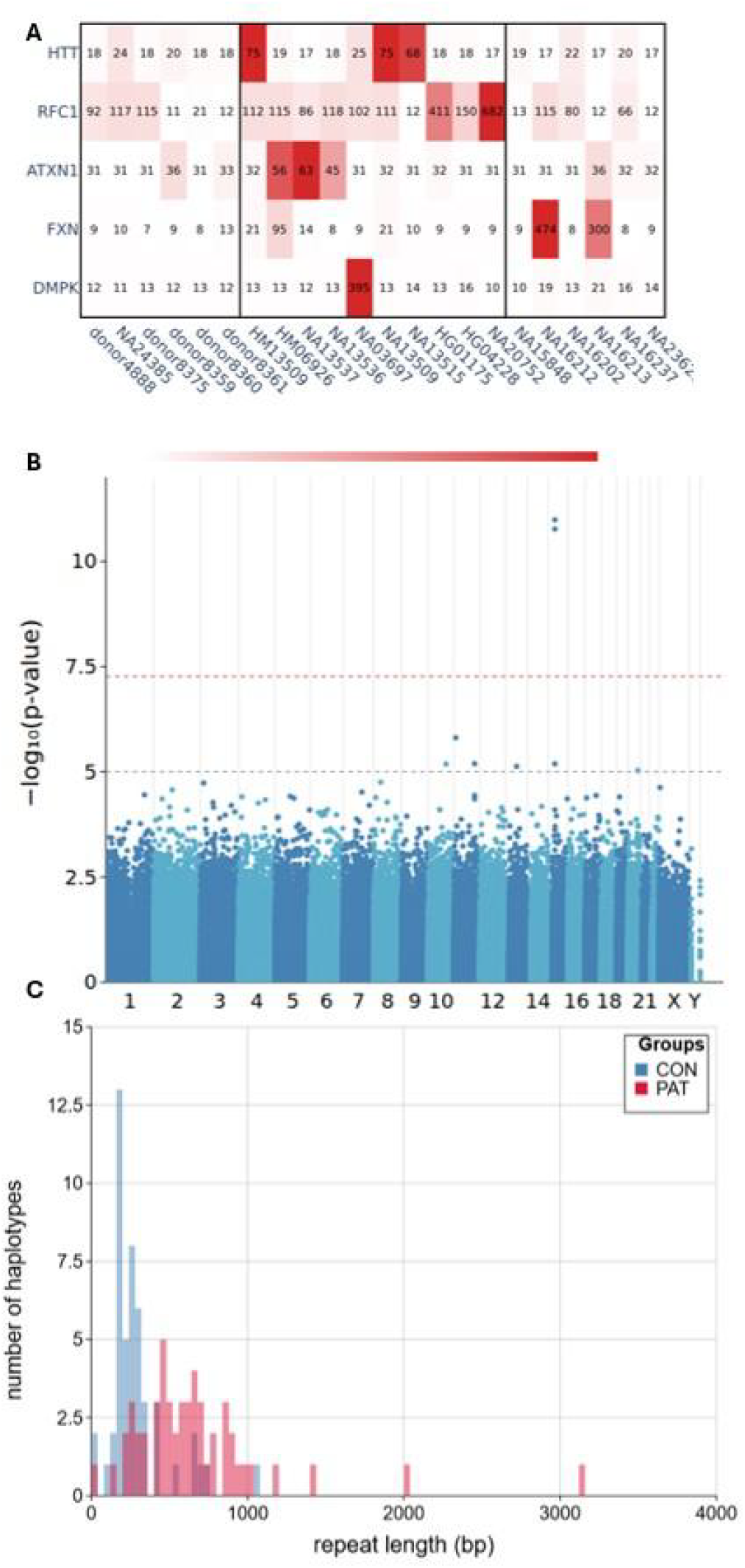
**A:** inquiSTR genotypes obtained based on public PacBio Puretarget data. The first block shows donors without repeat expansions; the second and third blocks contain known expanded alleles, with the third block containing *FXN* GAA expansion carriers, which remain a problematic composition for PacBio sequencers, leading to strongly underrepresented expanded alleles. **B:** Manhattan plot showing the result of inquiSTR association for the *GOLGA8A* expansion in aFTLD-U. **C:** Histogram of repeat lengths of the *GOLGA8A* expansion, showing longer fragment lengths in aFTLD-U patients.

## Discussion and conclusion

inquiSTR addresses a missing piece in population-scale tandem repeat research by enabling efficient and accurate length genotyping of millions of loci across hundreds of samples. Benchmarking on a single well-characterized sample demonstrates that inquiSTR is substantially faster and more memory-efficient than existing tools. Additionally, inquiSTR is at least as accurate as existing sequence-level genotypers, as inquiSTR genotypes are more similar to the GIAB truth genotypes. However, inquiSTR’s purpose is to complement, rather than replace, sequence-level genotypers such as TRGT and LongTR. The most practical workflow emerging from this work is a two-stage approach: inquiSTR is used to rapidly filter the full catalog to loci showing meaningful length variation, followed by a sequence-level genotyper for only that subset to also investigate the repeat’s sequence composition and motif variation, which is the main limitation of inquiSTR. This two-stage system substantially reduces computational burden without a meaningful loss of sensitivity. Beyond genotyping, inquiSTR offers several methods for follow-up of TR genotypes. We envision that inquiSTR’s modular structure will enable further extensions, such as recently developed methods for investigating population structure (Xia et al. 2026), beyond the modest separation in superpopulations achieved with PCA. Finally, genome-wide TR profiling is a crucial component in investigating missing heritability and causal mutations in human traits, complex disorders, and rare monogenic diseases, and these research questions are directly enabled by inquiSTR’s tailored subcommands for length-based association testing and length outlier detection. As long-read sequencing cohorts continue to grow, tools like inquiSTR make systematic TR profiling at the population scale not only feasible, but a routine.

## Methods

### Implementation

inquiSTR is a command-line toolkit for TR analysis from long-read sequencing data, designed around a TR length genotyping engine and an ecosystem of downstream analysis subcommands for population-scale studies. inquiSTR achieves computational efficiency through parallel processing and keeps the memory usage constrained through streaming algorithms. inquiSTR is implemented in Rust (Matsakis and Klock 2014) and depends majorly on rust-htslib, rust-bio, and noodles for handling BAM and CRAM files (Köster 2016; Macias 2026), rayon for parallelization (Matsakis 2024), and kuva for visualizations (Ferguson 2026).

### Repeat-length genotyping

The primary functionality of inquiSTR is TR length genotyping from aligned BAM or CRAM files through the `inquiSTR call` subcommand. The alignment files can be local or remote via HTTP/HTTPS/FTP/S3 URLs. inquiSTR scales efficiently to catalogs of millions of TR targets and accepts target regions through three mechanisms: (1) direct region specification via command-line string (e.g., `chr4:3074877-3074933`), (2) BED format files containing multiple loci, or (3) predefined preset catalogs. The preset system provides access to TR catalogs, with automated downloading on first use and local caching, and includes pathogenic disease-associated TRs from STRchive (75 loci) (Hiatt et al. 2024), forensic CODIS markers (20 loci) (Wang et al. 2022), and comprehensive genome-wide catalogs from Adotto (1.8M loci) (English et al. 2024) and TRExplorer (4.9M loci) (Weisburd et al. 2025). This approach can be further extended with other catalogs and genome build versions.

A performance optimization in inquiSTR is the locus batching system, which groups nearby TR targets within a configurable distance threshold (default 10 kb) for joint extraction of overlapping reads from the BAM/CRAM file, reducing I/O overhead and repeated fetches. Batches are further processed across multiple threads. Because performance depends on system architecture, input files, and system load, inquiSTR also includes an `optimize-call` subcommand that empirically determines the optimal combination of batch size and number of parallel threads.

The genotyping algorithm parses the CIGAR strings of reads overlapping TR loci to track insertions, deletions, and soft-clips, thereby determining the repeat-length difference relative to the reference genome. A genotype per haplotype is obtained by calculating the median across reads, using either phasing information (HP tags) from tools such as Longshot or WhatsHap (Edge and Bansal 2019; Martin et al. 2016), or by splitting unphased reads into two haplotypes based on their lengths. Spanning reads are prioritized for accuracy, and soft-clipped reads are added as needed to achieve sufficient support for genotyping the locus; a NaN genotype is returned if insufficient supporting reads are available (default: 3). When splitting reads into alleles, inquiSTR supports imbalanced allelic ratios, as is common in e.g., target capture data with long expanded alleles. The --haploid command-line argument specifies for which (sex) chromosomes splitting into two alleles should not be attempted. By default, the resulting genotypes are written in TSV format, sorted by chromosome and position, with two allele lengths (positive values indicate expansions, negative values indicate contractions), yielding a compact, human-readable, and easily parsable format. VCF output is also supported for compatibility with existing pipelines. The genotyping algorithm includes a filter for so-called foldback reads, an artifact in Oxford Nanopore data in which both the template and the complementary strands are sequenced consecutively (Heinz et al. 2026).

Alternatively, the `inquiSTR unmapped` subcommand profiles kmer frequencies from reads that were not mapped to the provided genome reference. The implementation computes all kmer sizes from 2 to a configurable maximum length (default 6), representing each kmer by its lexicographically smallest rotation to ensure consistent counting and optionally including the reverse complement or specifying a target kmer of interest using shorthand notations (e.g., quantify “(CAG)10”). Kmer frequencies are normalized to the total read count and output in TSV format compatible with downstream inquiSTR commands.

The `inquiSTR combine` subcommand merges individual results into a wide-format matrix for querying and statistical analysis. The combine command also supports incremental cohort building by adding new samples to an existing combined file. For cohort studies, the `inquiSTR batch` subcommand provides end-to-end processing from individual BAM/CRAM files to combined multi-sample results. This eliminates the need for users to manually orchestrate hundreds of separate commands. The batch command starts with a TSV file specifying samples and paths, processes samples in parallel, and can resume a crashed run.

### Downstream analysis subcommands

To follow up on TR genotypes, several subcommands are included in inquiSTR for visualization and statistical analysis. For compatibility with existing genotyping tools that generate VCF files, the `inquiSTR convert` subcommand can convert one or more VCF files into inquiSTR-compatible TSV files. The `inquiSTR query` subcommand provides rapid genotype lookup from combined files. The haplotypes are sorted by descending allele length to facilitate the identification of expanded alleles. TR lengths can be visualized either directly on the command line or using interactive HTML plots (Plotly Technologies Inc. 2015) with `inquiSTR plot` and `inquiSTR histogram`, enabling visual assessment of variation and comparison of lengths across groups (e.g., patients and control individuals). The `inquiSTR filter` subcommand enables locus prioritization and selection from individual or combined files using minimum TR length change, call rate, coefficient of variation, or genomic region. One major application we foresee is the use of inquiSTR in tandem with sequence-level genotypers, to initially perform length genotyping and then follow up on loci of interest, e.g., those with sufficient length variation or expansions, using a TR sequence-genotyping tool such as TRGT (Dolzhenko et al. 2024).

The `inquiSTR benchmark` subcommand enables comparison of inquiSTR calls against truth datasets in VCF or BED format (English et al. 2024), generating correlation plots and statistics, with options to filter by locus size, set a tolerance threshold, and output loci with the largest discrepancies. The user can specify which allele to use for comparison, either the longest (by default) or the shortest. This enables the evaluation of genotyping performance when adjusting parameters and the identification of challenging loci or systematic biases.

The `inquiSTR pca` subcommand performs principal component analysis on a combined file, to investigate (technical) outliers and population structure. The implementation offers three methods for aggregating haplotypes (max, min, or sum of allele lengths). The first two principal components are visualized in an interactive HTML plot (Plotly Technologies Inc. 2015), while additional components can be exported, for example, as covariates for association testing.

The `inquiSTR association` subcommand enables testing for differences in repeat length between groups (binary trait, e.g., case-control) or among all samples (continuous phenotype) by logistic and linear regression, respectively. Users specify a TR aggregation method (MEAN, MAX, or MIN for combining H1/H2 alleles), quality filters (call rate thresholds and minimum expansion lengths), and whether to include sample-level covariates in the model, while leveraging parallel processing across loci. The output includes effect sizes, p-values, and multiple testing correction using the Bonferroni method. The `inquiSTR outlier` subcommand identifies individual samples with TR lengths (or kmer frequencies) that are outliers relative to the cohort distribution, using either z-score thresholding (by default, ±3 standard deviations) or DBSCAN clustering. This is valuable for detecting pathogenic expansions when the number of carriers is insufficient to achieve statistical significance in an association study. The implementation enables focusing outlier analysis on a subset of the cohort (e.g., one or more patients) for rare disease analysis.

### Comparison of TR genotyping tools

The comparison between inquiSTR, TRGT, LongTR, and Straglr is based on data from the HG002 sample, sequenced on either an ONT PromethION (https://epi2me.nanoporetech.com/giab-2025.01/) or a PacBio Revio (https://downloads.pacbcloud.com/public/revio/2024Q4/WGS/GIAB_trio/HG002_rep1/m84039_241001_220042_s2.hifi_reads.bc2018.bam). The ONT dataset was downsampled using samtools (Li et al. 2009) to obtain coverage comparable to that of the PacBio data. For genotyping, the Adotto repeat catalog v1.2.1 (English et al. 2024) was used, except for the comparison of inquiSTR with Straglr, for which we created a smaller catalog by subsetting to chr21, removing homopolymers and loci with repeat motifs longer than 50 nucleotides, resulting in a set of 7596 repeat loci. All comparisons were performed on a local server running AlmaLinux 9.7 as the operating system, with AMD EPYC Genoa 9554P UP 64C/128T 3.1G CPUs and 16*64 DDR5, 4800 MHz RAM, and data read from SSDs. The comparison was executed using a snakemake workflow (Koster and Rahmann 2012), ensuring that no more than one tool is executed in parallel when benchmarking, running each tool/configuration with 5 replicates. VCF files were converted to inquiSTR format using inquiSTR convert and compared with the inquiSTR benchmark subcommand, using the longest allele at each locus for comparison. The tested tools are inquiSTR (v0.27.0), TRGT (v4.1.0), LongTR (v1.2), and Straglr (v1.5.6).

### Genotyping pathogenic TR loci from targeted sequencing

The ability of inquiSTR to genotype pathogenic TR loci was assessed using publicly available PacBio Revio sequencing data enriched for TR loci using the Puretarget method (see Data Availability section), and aligned with minimap2 (Li 2021). Because in target enrichment the allelic ratios between the expanded and reference alleles are typically skewed, and insufficient SNVs are available for phasing reads, the inquiSTR call command was run with --unphased and --imbalance 0.1 to improve sensitivity to longer expansions.

### Evaluating population structure

The performance of inquiSTR for investigating population structure (using PCA) was evaluated using public Oxford Nanopore Technologies PromethION from the 1000 Genomes Project (Noyvert et al. 2023), based on 20 individuals per superpopulation (self-reported geographical ancestry: AFR, AMR, EAS, EUR, and SAS). InquiSTR call was used to genotype a set of 171k polymorphic repeat loci (https://zenodo.org/records/7987365) (Dolzhenko et al. 2019) for the selection of individuals for PCA.

## Declarations

### Ethics approval and consent to participate

The ethics committee of the University Hospital Antwerp and the University Antwerp approved the study on GOLGA8A.

### Consent for publication

Not applicable

### Availability of data and materials

inquiSTR is distributed as precompiled static binaries compatible with Linux and macOS. Source code and extensive documentation are available under the MIT license at https://github.com/wdecoster/inquiSTR. The code is tested continuously and adheres to formatting and security standards, with automated dependency updates via GitHub Actions. The plots and comparisons presented in this manuscript are generated using the snakemake workflow and scripts in https://github.com/wdecoster/inquistr_paper.

Public PacBio sequencing data from pathogenic TR enriched with the Puretarget method was downloaded from https://downloads.pacbcloud.com/public/dataset/PureTarget2.0/PureTarget-Repeat2.0-Datasets/Repeat2.0_NanobindCoriell_48plex_RevioSRPQ/01-run-folder/1_A01/hifi_reads/

### Competing interests

WDC has received free consumables and travel reimbursement from Oxford Nanopore Technologies. WDC and RR are inventors on a patent filed concerning diagnostic applications of the GOLGA8A repeat expansion (WO/2026/062178). The remaining authors declare no competing interests.

### Funding

WDC receives a postdoctoral fellowship from the Flanders Fund for Scientific Research (FWO) 12ASR24N, and FK a postdoctoral fellowship from the Brein Instituut. This work was partly funded by the VIB (Flanders Institute for Biotechnology, Belgium) [RR, KS] and the University of Antwerp [RR, KS].

### Authors’ contributions

WDC developed inquiSTR, performed the benchmarking and comparisons, and wrote the manuscript. FK implemented the inquiSTR code to test the association between traits and repeat length. KS and RR oversaw and coordinated the project. All authors read and approved the final manuscript.

## Acknowledgements

Not applicable

## Notes

https://github.com/wdecoster/inquiSTR

