## Supplementary Figures for "inquiSTR: a toolkit for accurate and efficient population-scale tandem repeat genotyping and analysis"

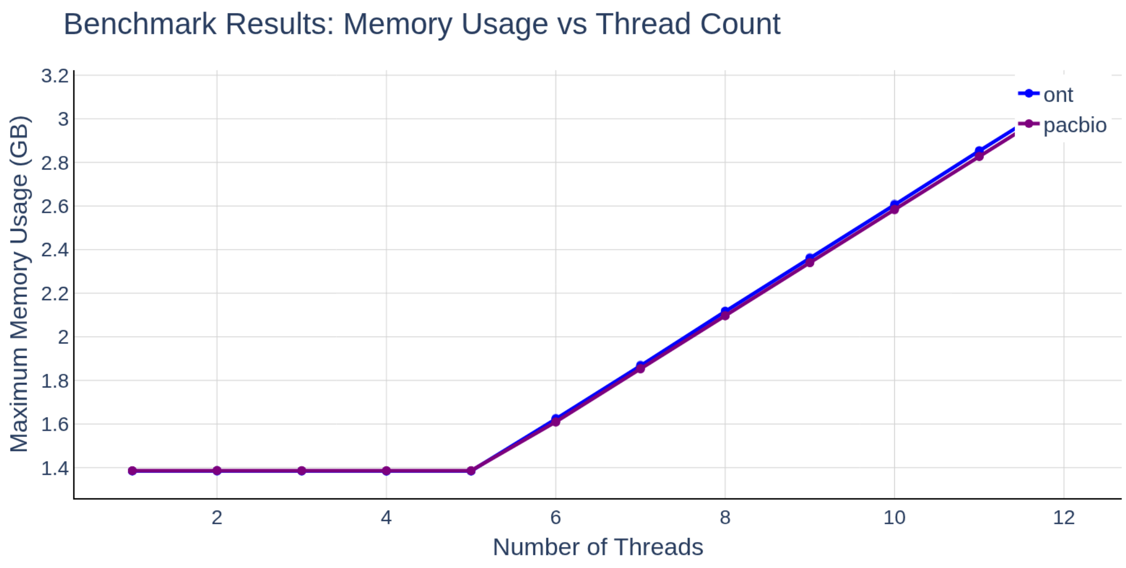


**Supplementary Figure 1**: memory use with increasing number of threads, accumulating linearly once more than 5 threads are used.

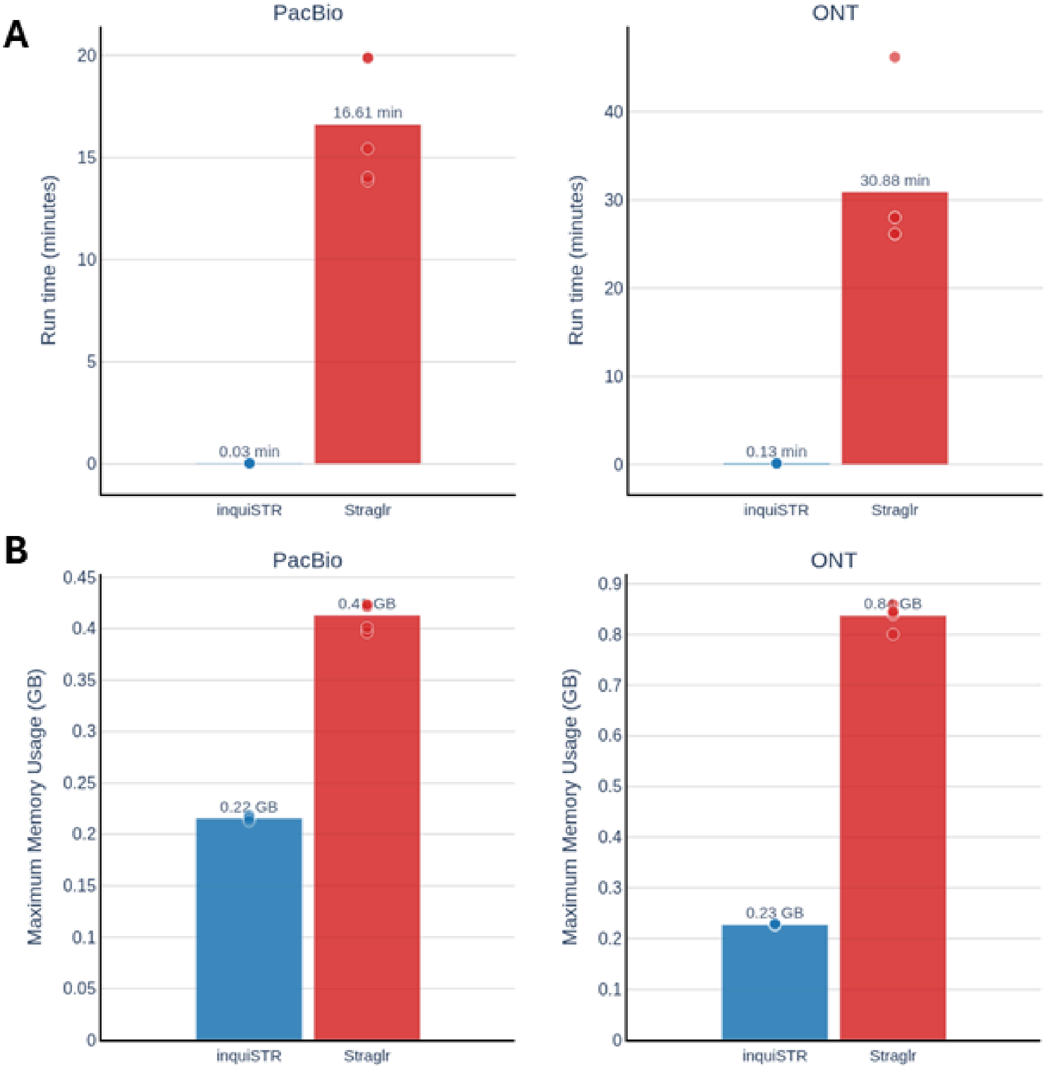


**Supplementary Figure 2**: Comparison of inquiSTR call genotyping with Straglr on a catalog of 7596 repeat loci. **A:** comparison of run time for the PacBio (left) and ONT (right) dataset. **B:** comparison of memory use for the PacBio (left) and ONT (right) dataset.

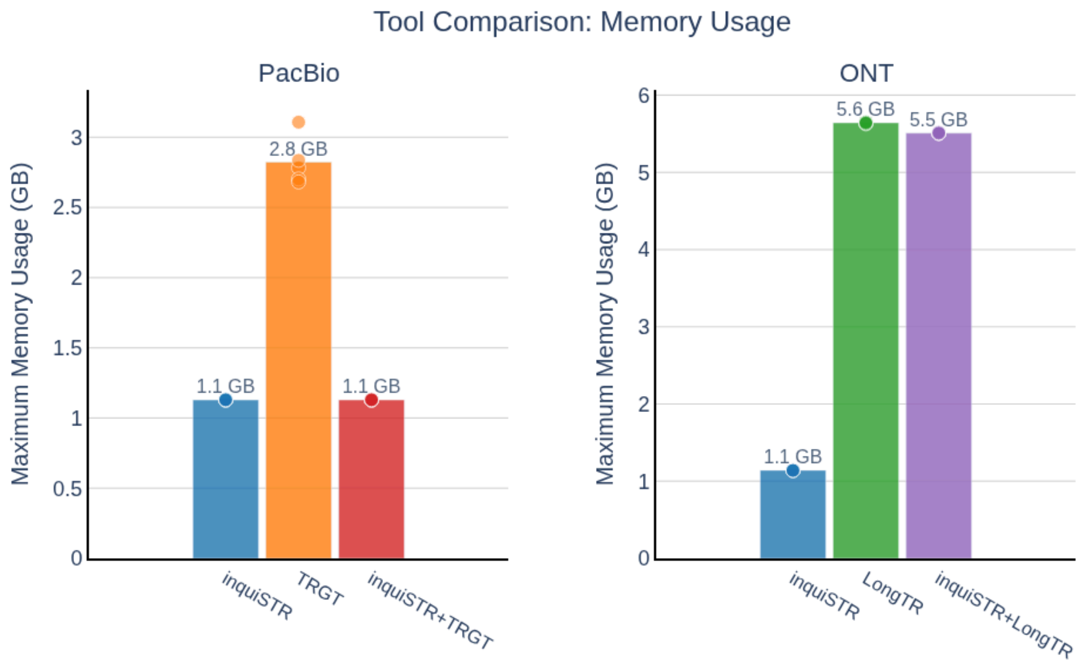


**Supplementary Figure 3:** comparison of memory usage of inquiSTR vs TRGT and LongTR on respectively PacBio and ONT data.

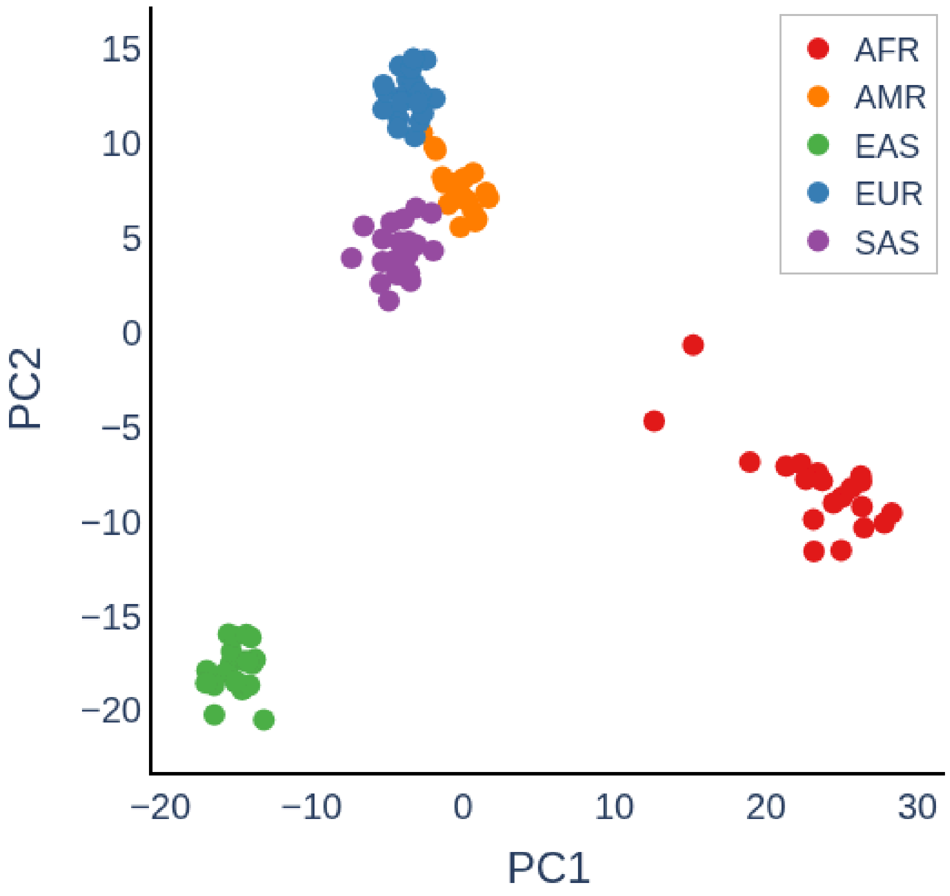


**Supplementary Figure 4:** Principal components analysis of 100 samples from the 1000 Genomes Project sequenced using ONT, colored by self-reported superpopulation.
